# *Lodhrasavam* modulates liver–adipose–pancreas axis to ameliorate MASLD: Observations from *In vitro* and *in vivo* models of hepatic and metabolic dysfunction

**DOI:** 10.1101/2023.07.05.547893

**Authors:** Sania Kouser, Subrahmanya Kumar, Chethala N Vishnuprasad

## Abstract

**Background:** Metabolic-dysfunction associated steatotic liver disease (MASLD) is a complex, multifactorial condition and a leading cause of liver-related morbidity worldwide. Due to its heterogeneous pathogenesis, conventional single-target therapies often show limited efficacy. Multicomponent *ayurveda* formulations offer a promising alternative through their multitargeted actions. This study explores the therapeutic potential and underlying mechanisms of *Lodhrasavam* (LS), a classical, polyherbal *ayurveda* formulation, in both in vitro and in vivo models of MASLD.

**Methods:** LS was evaluated for its anti-steatotic and anti-obesogenic potential using in vitro (HepG2, 3T3-L1) and in vivo (HF-HFD-induced rat) models. Antioxidant, lipase inhibition, cytotoxicity, anti-steatotic, and anti-adipogenic activities were assessed via DPPH, MTT, ORO/BODIPY staining, triglyceride quantification, ROS assay, and qPCR. Lipidomic profiling was done in HepG2 cells. Biochemical analyses (GLP-1, insulin, lipid panel), histopathology and OGTT further validated efficacy. Data were statistically analyzed by one-way ANOVA (*p*<0.05). Results/discussion: In HepG2 cells, LS reduced PA-induced lipid accumulation, ROS, triglycerides, and restored viability, while downregulating lipogenic genes (PPARγ, SREBP-1c, FASN). Lipidomics confirmed lipid modulation. In 3T3-L1 cells, LS suppressed adipogenesis and key adipogenic genes. In HFD-fed rats, LS reduced weight gain, hepatic steatosis, serum lipids, and improved liver histology. LS enhanced insulin and GLP-1 secretion, improved glucose tolerance, and restored pancreatic islet structure. These findings highlight the multitargeted potential of LS in ameliorating MASLD by modulating lipid metabolism, oxidative stress, inflammation, and glucose homeostasis.

**Conclusion:** *Lodhrasavam* ameliorates MASLD by modulating the liver–adipose–pancreas axis, improving lipid metabolism, reducing inflammation, and enhancing insulin and GLP-1 secretion through multitargeted mechanisms.

## Introduction

Non-alcoholic fatty liver disease (NAFLD) is a complex, multisystem disease, extending its effect on extrahepatic organs and regulatory pathways (Byrne and Targher, 2015). High-calorie diet and sedentary lifestyle are typical risk factors for NAFLD. However, many other factors including, but not limited to, type 2 diabetes, insulin resistance, obesity, hypertension, metabolic syndrome, genetic variations, alterations in gut permeability and microbiome are also playing a crucial role in its development and progression (Carr et al., 2016; Friedman et al., 2018; Younossi et al., 2018). It is one of the major contributors of liver disease worldwide with an estimated prevalence of 25% in the adult population (Younossi et al., 2019). In India its prevalence varies between 9% in rural and to 35% in urban populations (Duseja, 2010). When left untreated, NAFLD leads to deleterious liver physiology like steatohepatitis, fibrosis and cirrhosis that can further increase the risk of liver cancer and end-stage liver disease (Haas et al., 2016; Michelotti et al., 2013). In 2023, a Delphi consensus led by an international panel of experts resulted in the renaming of NAFLD to MASLD which is Metabolic-dysfunction associated steatotic liver disease. This globally supported transition marks a significant, non-stigmatizing shift in terminology and diagnostic criteria, aligning more accurately with underlying metabolic dysfunction. The change aims to enhance clinical clarity, facilitate improved patient identification, and reflect a more inclusive approach (Rinella et al., 2023).

Despite its high clinical and socioeconomic burden there are no approved therapeutics for MASLD and the current clinical management strategies are largely centered around controlling the predisposing factors like diabetes and obesity (Mundi et al., 2020). One of the reasons for this is the inadequacy of conventional single target-based drugs to address the complex etiopathology of MASLD as well as the limited knowledge we have on the systemic biology of this disease. Therefore, frontiers of biology started looking at innovative systemic strategies for addressing the multifactorial pathophysiology of MASLD. This is where a transdisciplinary approach of integrating biomedicine’s molecular perspective of MASLD with its systemic counterpart from traditional medicines like *Ayurveda* become relevant and can probably open up new opportunities for better holistic strategies for the prevention and management of MASLD.

One of the characteristics of MASLD is the accumulation of lipid droplets (LDs) in the hepatocytes leading to a condition referred to as hepatic steatosis, and it happens via flux of free fatty acids (FFAs) from adipose tissue due to increased lipolysis, *de novo lipogenesis* (DNL), excess dietary fat intake, decreased export of triglycerides and reduced beta-oxidation (Alkhouri et al., 2009; Schulze and McNiven, 2019). Although *Ayurveda* has an epistemologically and ontologically different understanding of MASLD from that of biomedicine, both schools emphasize the involvement of metabolic imbalances and the role of diabetes (*Prameha*) and obesity (*Sthoulya*) as predisposing factors in MASLD. Previous studies from our laboratory by Butala et al., 2017 reported the anti-diabetic and anti-obesity effects of *Lodhrasavam*, a polyherbal formulation in *Ayurveda* prescribed for obese-diabetic patients. According to Ayurvedic pharmacology principles, the key herbal ingredients of this formulation belong to the *Lodhradi-gana*, a group of plants having *medo-hara* (anti-obesity) and *kapha-hara* properties and both are essential for the management of glucose and lipid homeostasis in the body. Given the potent anti-obesity and anti-lipidemic properties of *Lodhrasavam*, along with its diverse herbal composition, the present study aimed to evaluate its multisystemic efficacy in managing MASLD by targeting hepatic steatosis, adipogenesis, and the regulation of metabolic hormones.

## Materials and methods

### Chemicals and Reagents

HepG2 human hepatoma cell line (NCCS, Pune, India); Dulbecco’s modified Eagle’s medium (DMEM) (GIBCO, Grand Island, New York); fetal bovine serum (FBS) (GIBCO, Grand Island, New York); penicillin/streptomycin (GIBCO, Grand Island, New York); fatty acid free bovine serum albumin (BSA) (Genei, Bangalore, India); sodium palmitate (PA) (Sigma-Aldrich, St. Louis, MO, USA); 3-(4,5-dimethylthiazol-2-yl)-2,5-diphenyltetrazolium bromide (MTT) (SRL, India); triglyceride kit (BeneSphera, Avantor, USA); 2,2-Diphenyl-1-picrylhydrazyl (DPPH) (Sigma-Aldrich, St. Louis, MO, USA); Ascorbic Acid (Sigma-Aldrich, St. Louis, MO, USA); Folin – Ciocalteu reagent (Sigma-Aldrich, St. Louis, MO, USA); gallic acid (Sigma-Aldrich, St. Louis, MO, USA); BODIPY 493/503 (Cayman chemical 25892); Hoechst 33342 (Cayman chemical 15547); Oil-Red-O stain (Sigma-Aldrich, St. Louis, MO, USA); trizol reagent (Takara Bio, Shiga, Japan); PrimeScript 1st strand cDNA Synthesis Kit (Takara Bio, Shiga, Japan); PrimeScript RT-PCR Kit (Takara Bio, Shiga, Japan)

### Determination of total tannin content

LS was estimated for total tannins following standard Folin - Ciocalteu method (Ainsworth and Gillespie, 2007). Briefly, 10μL of LS with 40μL of water, 50μL FC reagent (previously diluted with water 1:10 v/v) and 100μL of 3.5% Na2CO3 was incubated at room temperature for 30 min. A set of gallic acid standards (50, 25, 12.5, 6.25, 3.125μg/mL) were prepared in the same manner. The absorbance was measured at 760 nm using plate reader (xMark Microplate Spectrophotometer, BioRad, USA). The experimental concentrations of test samples were expressed as μg of gallic acid equivalent tannin (GAE)/mL of sample.

### Determination of DPPH radical scavenging activity

Free radical scavenging ability of different concentrations of LS was tested by DPPH radical scavenging assay (Rahman et al., 2015). DPPH produces violet/purple color in ethanol solution and fades to shades of yellow color in the presence of antioxidants. A solution of 0.1 mM DPPH in ethanol was prepared, and 2.4 mL of this solution was mixed with 1.6 mL of LS in ethanol to get a concentration gradient (128, 64, 32, 16, 8, 4, 2μg GAE/ml). The reaction mixture was vortexed thoroughly and left in the dark at room temperature (RT) for 30 min. The absorbance of the mixture was measured spectrophotometrically at 517 nm. Ascorbic acid was used as reference. Percentage DPPH radical scavenging activity was calculated by the following equation: % DPPH radical scavenging activity= ((A_0_− A_1_)/A_0_) ×100; where A_0_ is the absorbance of the control, and A_1_ is the absorbance of LS or the standard. The % of inhibition was plotted against concentration, and from the graph IC_50_ was calculated. The experiment was performed in triplicates and repeated three times. Ascorbic acid Equivalent Antioxidant Capacity (AEAC) was calculated using the formula: IC_50_ (AA) /IC_50_ (sample). Where, IC_50_(AA) is IC_50_ of standard (i.e., Ascorbic acid) and IC_50_(sample) is IC_50_ of sample.

### Pancreatic lipase inhibition

Different concentrations of LS were aliquoted and made up to 50µl using 50mM Sodium Phosphate buffer (pH – 8). To this, 150µL of lipase enzyme mixture (10mg/mL) was added and incubated for 30 mins at 37°C. After incubation, 100 µL of p-nitrophenylbutyrate (1mM) and incubated at room temperature for 20 mins. The absorbance was read at 405 nm using spectrophotometer.

### Cell Culture and establishment of palmitic acid induced MASLD model

HepG2 cells were cultured in DMEM containing 10% FBS and 1% penicillin/streptomycin in a humidified atmosphere containing 5% CO2 at 37°C. The cells were seeded in multi-well plates and grown for 48 hrs. The medium was changed to DMEM (serum free) containing 5% fatty-acid-free bovine serum albumin (BSA) with or without 1mM palmitic acid (PA) and grown for 24 h. Further, the cells were treated with various concentrations of LS in the same BSA-medium and incubated for 24 h. The changes in lipid droplet formation were observed and quantified using various methods as described in the below sections.

### Cytotoxicity assay

The cytotoxic effects of various concentrations of LS in HepG2 cells were studied using MTT assay, which is based on the reduction of a tetrazolium salt by mitochondrial dehydrogenase in viable cells. HepG2 cells were seeded in 96-well plates at the density of 1.5 × 10^4^ cells/well and cultured in 100μL of DMEM containing 10% FBS overnight. The medium was then discarded and was replaced with 128, 64, 32, 16, 8, 4, 2, 0μg GAE/ml LS diluted in the culture medium, followed by incubation for a further 24 h at 37°C. 500μg/ml of MTT working solution were then added to each well followed by incubation for 4 h at 37°C. The violet formazan crystal in each well was dissolved in DMSO and the absorbance of each well was measured at 570 nm using a microplate reader (xMark Microplate Spectrophotometer, BioRad, USA). HepG2 cells were treated with 1mM PA in 5% BSA media for 24 h at 37°C, followed by treatment with 128, 64, 32, 16, 8, 4, 0μg GAE/ml LS for 24 h at 37°C. Further, to assess they cytotoxicity effect MTT assay was performed as described.

### BODIPY 493/503 staining of lipid droplets

HepG2 cells were seeded in 35mm confocal dishes, treated with 1 mM PA for 24 h, and then cultured with 32μg GAE/ml LS for 24 h at 37°C. Cells were stained with 10μM BODIPY 493/503 dye and 10μg/ml Hoechst dye and incubated for 30 mins at 37°C (Qiu and Simon, 2016). Cells were washed with PBS, visualized under the LSM880 confocal live cell imaging system, (Carls Ziess, Germany) and images were taken at 40x magnification.

### Oil Red O (ORO) staining of intracellular lipid accumulation

HepG2 cells were seeded in six-well plates (1.2×10^6^ cells/well), treated with 1 mM PA for 24 h, and then cultured with 32μg GAE/ml LS for 24 h at 37°C. Cells were washed twice with PBS, and fixed with 4% formaldehyde for 30 mins. After washing, cells were stained for 20 min in freshly diluted ORO solution (0.5% ORO in isopropanol diluted to 3:2 with H2O, and filtered) at 37 °C. After staining, the cells were thoroughly washed with double distilled water to remove the unbound staining solution and images of cells stained with Oil Red were captured using phase contrast microscope (Olympus-IX-Olympus America Inc, USA). To quantify the ORO contents, isopropanol was added to each well and the optical density of the samples at a wavelength of 450 nm was measured using a microplate reader (xMark Microplate Spectrophotometer, BioRad, USA) (Mehlem et al., 2013).

### Triglyceride (TG) estimation

To analyze the content of cellular triglycerides, the cells were washed with PBS, scraped into 250μl 0.1% PBST and pulse sonicated for 45 % amplitude for 45 seconds. The lysates were assayed for their total triglyceride content using assay kit and cellular protein was estimated using the Bradford method. The results were expressed as µg of triglyceride per mg of cellular protein.

### Measurement of cellular ROS content

HepG2 cells were seeded in 12 well plates, treated with 1 mM PA for 24 h, and then cultured with 32μg GAE/ml LS for 24 h at 37°C. The cells were incubated with 10μM fluorescent probe 2,7-dichlorodihydrofluorescein diacetate (DCDHF-DA) prepared in serum free DMEM media for 30 mins in dark at 37°C. After the incubation period cell were washed thrice with ice cold PBS and the fluorescence intensity was measured at Excitation: 485 and Emission: 530 using Synergy H1, Biotek microplate reader. The fluorescence intensity values were normalized with the protein content which was measured by the bradford method.

### Quantification real-time PCR

Total RNA was isolated from treated HepG2 cells using trizol reagent according to the manufacturer’s instructions, and the absorbances of the extracted RNAs at 260 nm and 280 nm were determined spectrophotometrically (Nanodrop, Thermofisher Scientific). RNA samples with purity ratios (A260/A280) between 1.8 and 2.0 were used for synthesizing single-strand cDNA by means of PrimeScript 1st strand cDNA Synthesis Kit for quantitative real-time polymerase chain reaction (qPCR). Gene-specific primers were designed (Table 3) (Bioserve, Hyderabad, India) and qPCR was performed to detect the relative mRNA expression levels using PrimeScript RT-PCR Kit as described in the protocol. To determine the specificity of the amplification, melting curve analysis was performed for all final PCR products. Relative changes in gene expression were calculated and expressed as fold changes using the relative quantification (ΔΔCt) method. The relative abundance of each transcript was normalized to that of GAPDH. Each sample was run in triplicate.

**Table 1.**
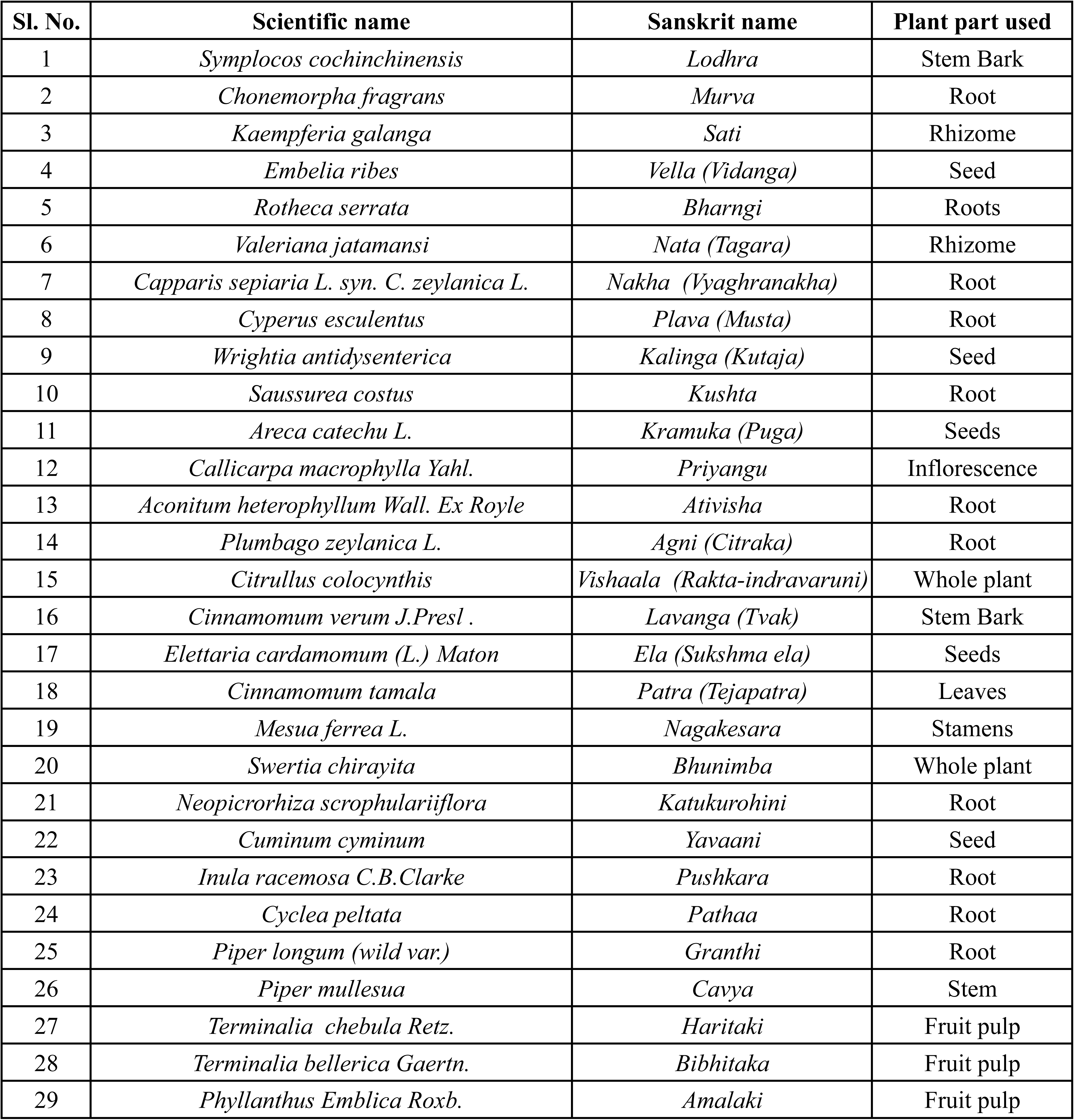
**List of plant ingredients present in *Lodhrasavam***

| Sl. No. | Scientific name | Sanskrit name | Plant part used |
| --- | --- | --- | --- |
| 1 | *Symplocos cochinchinensis* | *Lodhra* | Stem Bark |
| 2 | *Chonemorpha fragrans* | *Murva* | Root |
| 3 | *Kaempferia galanga* | *Sati* | Rhizome |
| 4 | *Embelia ribes* | *Vella (Vidanga)* | Seed |
| 5 | *Rothea serrata* | *Bharngi* | Roots |
| 6 | *Valeriana jatamansi* | *Nata (Tagara)* | Rhizome |
| 7 | *Capparis sepiaria* L. syn. *C. zeylanica* L. | *Nakha (Vyaghranakha)* | Root |
| 8 | *Cyperus esculentus* | *Plava (Musta)* | Root |
| 9 | *Wrightia antidysenterica* | *Kalinga (Kutaja)* | Seed |
| 10 | *Saussurea costus* | *Kushta* | Root |
| 11 | *Areca catechu* L. | *Kramuka (Puga)* | Seeds |
| 12 | *Callicarpa macrophylla* Yahl. | *Priyangu* | Inflorescence |
| 13 | *Aconitum heterophyllum* Wall. Ex Royle | *Ativisha* | Root |
| 14 | *Plumbago zeylanica* L. | *Agni (Citraka)* | Root |
| 15 | *Citrullus colocynthis* | *Vishaala (Rakta-indravaruni)* | Whole plant |
| 16 | *Cinnamomum verum* J.Presl . | *Lavanga (Tvak)* | Stem Bark |
| 17 | *Elettaria cardamomum* (L.) Maton | *Ela (Sukshma ela)* | Seeds |
| 18 | *Cinnamomum tamala* | *Patra (Tejapatra)* | Leaves |
| 19 | *Mesua ferrea* L. | *Nagakesara* | Stamens |
| 20 | *Swertia chirayita* | *Bhunimba* | Whole plant |
| 21 | *Neopicrorhiza scrophulariiflora* | *Katukurohini* | Root |
| 22 | *Cuminum cyminum* | *Yavaani* | Seed |
| 23 | *Inula racemosa* C.B.Clarke | *Pushkara* | Root |
| 24 | *Cyclea peltata* | *Pathaa* | Root |
| 25 | *Piper longum* (wild var.) | *Granthi* | Root |
| 26 | *Piper mullesua* | *Cavya* | Stem |
| 27 | *Terminalia chebula* Retz. | *Haritaki* | Fruit pulp |
| 28 | *Terminalia bellerica* Gaertn. | *Bibhitaka* | Fruit pulp |
| 29 | *Phyllanthus Emblica* Roxb. | *Amalaki* | Fruit pulp |

**Table 2.** Experimental and clinical effect of the individual plant ingredients in *Lodhrasavam* on NAFLD.

| Plant | Experimental/ Clinical Evidence |  |  |  |  |  |  |
| --- | --- | --- | --- | --- | --- | --- | --- |
|  | Model/ Study design | Induction | Extract/molecule | Biochemical parameters | Molecular markers | Clinical Summary | Reference |
| *Areca catechu* | Male C57BL/6N | 30% Lard, 1% cholesterol, 4% sugar water | Areca nut extract, areca nut polyphenol | ↓ Total liver lipids, serum levels of GPT and GOT, Cholesterol, non-HDL | ↑ phospho-AMPK, ↓ SREBP2, HMGCR |  | Yi et al., 2023 |
|  | Total subjects: 5967; Healthy individuals: 4052 and NAFLD individuals: 1915 |  | Betel nut chewing |  |  | In subjects with NAFLD, betel nut chewing, even ex-chewing, was associated with a higher risk of liver fibrosis | Chou et al., 2021 |
|  | Total subjects: 9221; Healthy individuals: 6981 and NAFLD individuals: 2240 |  | Betel quid chewing |  |  | Betel quid chewing was associated with significant liver fibrosis in subjects with MetS, but not in subjects without. | Chou et al., 2022 |
| *Cinnamomum verum J.Presl. (Cinnamomum zeylanicum Blume)* | Randomized double blinded, placebo-controlled trial; 45 NAFLD patients (23 patients in the intervention group, 22 patients in the control group) |  | Cinnamon capsule/ wheat flour capsule (placebo)- 2 capsules daily for 12 weeks | ↓ HOMA, FBS, AST, ALT, GGT, hs-CRP, Cholesterol, Triglyceride |  | The study suggests that taking 1500 mg cinnamon daily may be effective in improving NAFLD characteristics. | Askari et al., 2014 |
|  | Male Wister rats | HFD (43% Fat) | Cinnamon bark powder (50mg/Kg, 100mg/Kg) | ↓ Body weight, liver weight, FBG, HOMA-IR, AST, ALT, LDL, Total cholesterol, total triglyceride; ↑ HDL, SOD, Catalase, GPx | ↑ NRF2, CPT-1, CD-36, PPARA; ↓ NFKB, TNFA, SREBP-1c |  | Li et al., 2022 |
| | Male Swiss mice | 19.8% Lard, 19.8% corn oil | Gold nanoparticles with C. verum bioactives | ↓ Liver triglycerides, plasma triglycerides, HOMA-IR | ↓ IL-1β, IL-6, INF-γ, MCP-1 | | Sharma et al., 2022 |
| *Elettaria cardamomum (L.) Maton* | A double-blind randomized clinical trial; 87 subjects (cardamom n = 43; placebo n = 44) |  | Two 500-mg capsules (cardamom/ toast powder), three times/day, taken with meals for 3 months, follow-up monthly | ↑ irisin, HDL-c, and QUICKI; ↓ FBI, TG, LDL-c, HOMA-IR, and the grade of fatty liver |  | GC supplement improved the grade of fatty liver, serum glucose indices, lipids, and irisin level among overweight or obese NAFLD patients. | Daneshi-Maskooni et al., 2019 |
|  | Wistar strain | Dexamethasone (8 mg/kg) induced | cardamom powder suspension in 2% gum acacia 1 g/kg/10 ml | ↓ PPBS, FBS, Cholesterol, Triglyceride; ↑ HDL |  |  | Nitasha Bhat et al., 2015 |
| *Andrographis paniculata* (Burm.f.) Wall. ex Nees | HepG2; Male Wistar rats | 1mM Palmitate-oleate mixture; HFD- 20% lard | Andrographolide (AN), Isoandrographolide (IAN) and 3,19-acetonylidene andrographolide (ANA) | ↓ AST, ALT, ALP, steatosis, inflammation, ballooning, serum triglycerides, serum total cholesterol; reduced lipid droplets and triglycerides in cells |  |  | Toppo et al., 2017 |
| | Male C57BL/6J mice | HFHC diet - 25% lard, 0.5% cholesterol | 0.05% and 0.1% of 14-Deoxy-11,12-didehydroandrographolide (deAND), a diterpenoid in *Andrographis paniculata* | ↓ ALT, AST, hepatic cholesterol level, hs-CRP, | ↓ caspase 3/pro-caspase 3 ratio, TNFα, IL-1β, caspase 1, NLRP3, ↑ NRF2, GSH, HO-1 | | Liu et al., 2020 |
| | HepG2, Wistar rats | Palmitate-oleate mixture; HFD | 12.5–50 μM of andrographolide (AN), isoandrographolide (IAN), isoandrographolide diacetate (IAN-ACT) isoandrographolide 19 propionate (IAN-19P), isoandrographolide dipropionate (IAN-DP), isoandrographolide acetone (IAN-ACN) and isoandrographolide-19-benzoate (IAN-BEN); AN, IAN and IAN-19P were fixed as 25 mg/kg b.w of rat | ↓ ALT, AST, LDH leakage, intracellular triglycerides and lipid droplet accumulation (IAN-19P showed better results); ↓ ALT, AST, LDH, GGT, ACP, Cholesterol, Triglyceride, Ballonong, steatosis and inflammation | ↓ TNFα, SREBP-1c, PPAR-γ; ↑ PPAR-α, CPT-1, FXR | | Toppo et al., 2021 |
| | Male C57BL/6 J mice. LO2 cells | HFD; 0.5mM Oleic acid | Andro-L (50 mg/kg/day), Andro-M (100 mg/kg/day), and Andro-H (200 mg/kg/day); 20μM Andro | ↓ FBG, HbA1c, OGTT, triglyceride, total cholesterol; ↓ lipid accumulation, | ↓ SREBP-1c, PPAR-γ, LDLR, FATP1, FATP2, FATP3, FATP4, FATP5, FAS, SCD1, GPAT, DGAT1; ↑ PPAR-β, CPT-1, HSL | | Ran et al., 2023 |
| *Picrorhiza kurroa* Royle ex Benth. | Male Wistar rats | 30% HFD (Amul butter) | P. kurroa rhizomes ethyl-alcohol:water extract (200 mg/kg - 400 mg/kg) | ↓ liver lipid content, serum cholesterol, serum triglycerides, ALT, ALP |  |  | Shetty et al., 2010 |
| | HepG2 | 1mM Palmitate-oleate mixture | Picroside II (10μM) | ↓ lipid accumulation, ROS, ↑ GSH, ATP | ↑ MnSOD, Catalase, cytochrome c | | Dhami-Shah et al., 2021 |
|  | Male Wistar albino rats | Hz (Hydrazine sulphate) (50 mg/kg) | Picoliv (PIC) (50 mg/kg) | ↓ ALT, AST, bilirubin, cholesterol level, triglyceride, phospholipid, total lipid, free fatty acid |  |  | Vivekanandan et al., 2007 |
| | HepG2 | 500μM Palmitate-oleate mixture | Picroside II (10μM) | ↓ lipid accumulation | ↓ SREBP-1c, FATP5, SCD1, FOXO1, PEPCK | | Dhami-Shah et al., 2018 |
| *Cyperus rotundus* L. | male C57BL/6 mice; Mice primary hepatocytes; HepG2 | HSD diet (65% sucrose); | Hexane fraction of *Cyperus rotundus* rhizome | ↓ hepatic triglyceride, serum AST; ↑ HDL | ↓ SREBP-1c, SCD1, GPAT, FAS, ACC1 |  | Oh et al., 2015 |
| *Phyllanthus Emblica* Roxb. | male C57BL/6J mice | CDAHFD (choline-deficient, L-amino acid-defined) | Aqueous extract of *Phyllanthus Emblica* (0.9, 1.8, 3.6 g of crude drug/kg) | ↓ ALT, AST, triglyceride, LDL, lipid droplet; ↑ HDL |  |  | Luo et al., 2022 |
| | Male C57BL/6JNarl mice | MCD diet | Water extract of Phylanthus Emblica (125, 250, 500 mg/kg b.w.) | ↓ ALT, AST; ↑ catalase, GPx, GST, GRd | ↓ CYP2E1, TNF-α, IL-1β | | Tung et al., 2018 |
| | Male Sprague–Dawley rats | HFD (40% beef tallow) | Water extract of Phylanthus Emblica (125, 250, 500 mg/kg b.w.) | ↓ ALT, AST, LDL; ↑ HDL, GST, Catalase, GPx, | ↑ PPAR-α, adiponectin, leptin; ↓ SREBP-1c, LXR-α | | Huang et al., 2017 |
| | Female Wistar rats | Ovariectomized; High fructose diet (60%) | Ethyl acetate extract of amla (100 mg/kg body weight/day) | ↓ ALT, AST, Liver weight; | ↑ PPAR-α, LXR, Insig-2, ABCA1, LDLR; | | Koshy et al., 2015 |

**Table 3:**
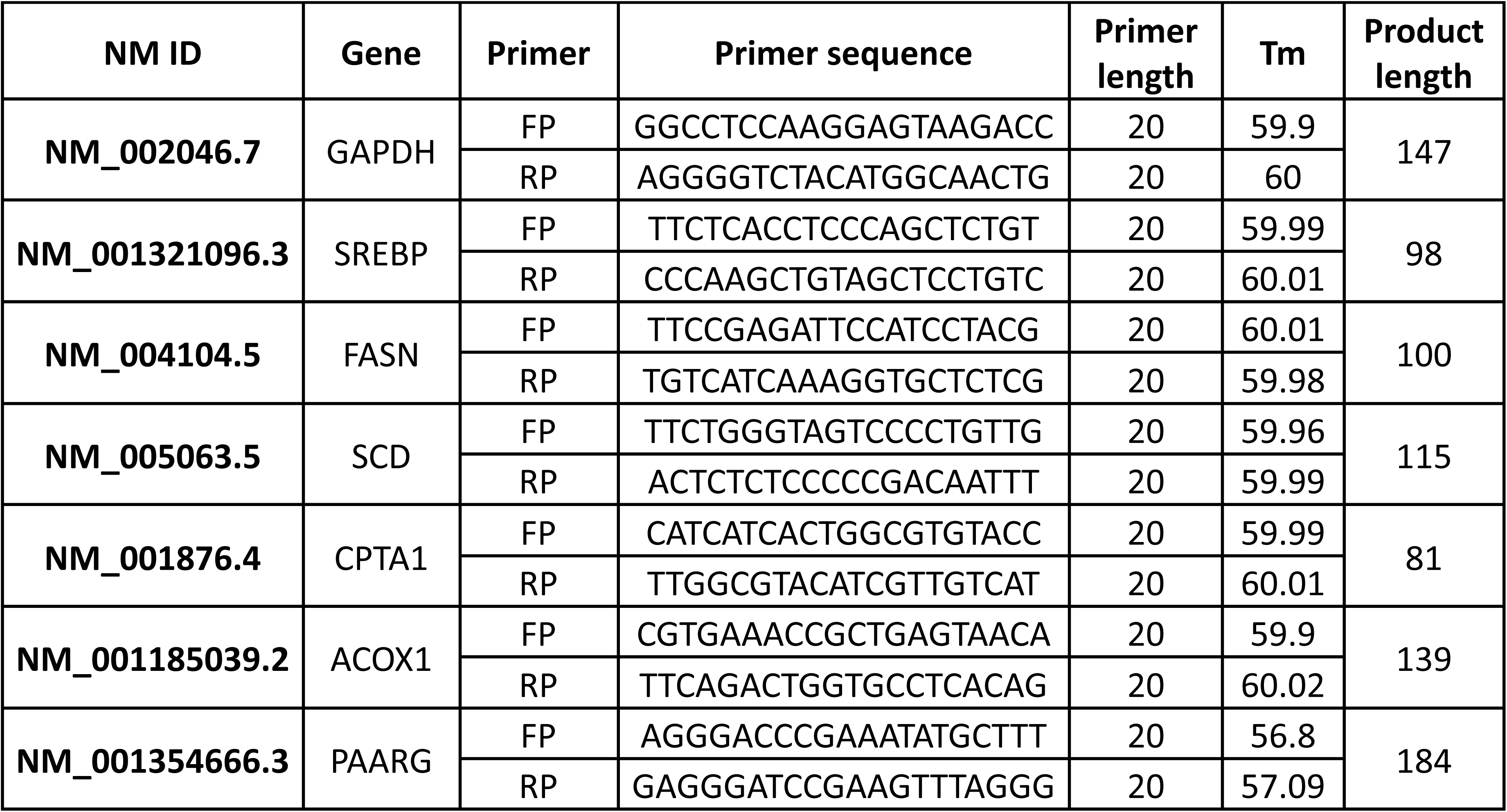
List of primers.

### Lipidomic analysis

The cells were seeded and were allowed to grow for 48hrs; they were then cultured with 5% fatty-acid-free bovine serum albumin (BSA) (Genei, Bangalore, India) in serum free DMEM (5% BSA media) media or palmitic acid (PA) (Sigma-Aldrich, St. Louis, MO, USA) in 5% BSA media for 24 h. Further, the cells were either treated with 5% BSA media, or treated with LS for 24 h. After which, cells were washed thrice with ice cold PBS and lipids were extracted following Bligh and Dyer method with slight modification. The cell pellet was spiked with 10μl of Internal standard (SPLASH^®^ LIPIDOMIX^®^ Mass Spec Standard, Avanti polar lipids, Merck) and ice cold 200μl of methanol, 200μl chloroform and 200μl 0.8% KCl was added to the pellet. The sample was pulse sonicated for 1 min followed by incubation at 37°c for 10 mins. The sample was then centrifuged at 10,000 rpm for 10 mins at 4°c, the lower organic phase was collected and dried in a speed vac for half an hour. The pellet was reconstituted in 100μl of methanol and 5μl of the sample was injected for untargeted lipidomic analysis in Orbitrap mass spectrometer. Lipidomic analysis was done using MS-DIAL software and Metaboanalyst 6.0. Aligned data obtained from MS-DIAL was subjected to statistical analysis in Metaboanalyst. Abundance data was filtered using variance filter of IQR 25% and mean intensity was used as abundance filter. Data was normalized using log_10_ data transformation and auto scaling was used for data scaling. For all the analysis p value < 0.05 was used. For pathway analysis BioPAN database was used.

### Anti-adipogenic effect of LS on 3T3-L1 cells

The 3T3-L1 fibroblasts were purchased from the National Centre for Cell Sciences, Pune, India. The cells were maintained in DMEM containing 1% penicillin and streptomycin supplemented with 10% FBS at 37°C in a 5% CO_2_ humidified atmosphere. The 3T3-L1 fibroblasts were differentiated by incubating two days post-confluent (day-0) plates in differentiation induction medium containing 500μM IBMX, 250nM dexamethasone and 50nM insulin (MDI). On day-3, the medium was replaced with insulin media for 2 days and subsequently the cells were maintained in fresh culture media till they attain complete adipocyte morphology. To check the anti-adipogenic effect of LS, 32µg of GAE/mL LS was added along with MDI and incubated. On day-7 fully differentiated adipocytes were used for Oil Red O staining, triglyceride assay and qRT-PCR.

### Animal model

All *in-vivo* experiments were carried out in Acharya & BM Reddy college of Pharmacy, Bengaluru, India. Male Sprague Dawley rats (SD rats) weighing 180-200 g were procured and maintained in Acharya & BM Reddy college of Pharmacy animal housing facility. Animals were maintained under controlled environmental conditions (temperature 21-25°C and normal humidity conditions). The animals were fed with commercial HFHFD procured from VRK nutritional solutions, Pune, India. The study protocol was approved by institutional animal ethics committee (IAEC/ABMRCP/2023-2024/4).

### Acute Toxicity studies

Based on the limit test standard of the Organization for Economic Cooperation and Development (OECD) No. 425 Guideline described in the reference for acute toxicity testing of drugs in rodents, 2 female SD rats were administered orally with 2mL/Kg of LS (Kifle and Belayneh, 2020). After 24hrs of observation for physical and behavioral change, the animals were further administered with a higher dose of 5mL/Kg of LS and observed for 14 days for any sign of toxicity. Formulation was well tolerated and showed no fatality or no signs of any abnormal behavior. For administration of formulation in MASLD condition, the doses were calculated according to the conversion table based on surface area as rat equivalent volume of adult human dose, derived by multiplying the adult human dose by 0.018 per 200 g of body weight of rat (Ghosh MN, 2015). The adult human dose of formulations was considered to be 30 ml as high dose and 15 ml as low dose after discussion with the *Ayurveda* physicians. Formulation was administered by oral gavage.

### Induction of MASLD

Male SD rats weighing 150±180g were randomly divided into two nutritional groups: a standard diet (control group) and a high-fat-high-fructose diet (HFHFD - 60% energy from fat (Palm oil), 2% Cholesterol, 20% from carbohydrate (Fructose), and 20% from protein; Total energy 5200 kcal/kg). Before the HFHFD feeding, animals were weighed and their initial blood glucose level were recorded. The animals were fed with HFHFD and daily food intake was recorded. Animals were weighed weekly to evaluate the weight gain. After 90 days of HFHFD diet, experimental rats were administered with the respective drugs with an oral gavage for a period of 30 days. The high fat diet was continued throughout the formulations’ treatment for the respective groups. Pioglitazone (PIOZ) was used as a positive control in the study.

### Oral Glucose Tolerance Test

Briefly, on the day of OGTT, animals were fasted for 3hrs, following that the animals were administered with 2 gm/Kg of glucose orally. Blood samples were collected from the caudal vein, by means of a small incision at the end of the tail, at 0, 30, 60 and 120 min after glucose administration. Blood Glucose Level (BGL) was estimated by the enzymatic glucose oxidase method using a commercial glucometer (Accucheck Active glucometer and glucose strips). The results were expressed as the integrated area under the curve for glucose (AUC glucose), which was calculated using MS excel.

### Biochemical parameter assessment

At the end of the experimental day, blood was collected from the retro-orbital sinus and centrifuged at 3500 rpm for 10 min at 4°C. The supernatant was obtained for GLP-1 measurement. The plasma samples collected from the animals were assayed for GLP-1 secretion using Raybiotech GLP-1 kit, (catalogue no: EIA-GLP1) and Insulin secretion using Krishgen Elisa kit. Serum was separated and analyzed spectrophotometrically for triglyceride, cholesterol, HDL and LDL levels using the standard assay kits.

### Histopathological analysis

At the end of the experiments, the liver, adipose and pancreas in each animal group were immediately removed and fixed in 10 % buffer saturated formalin. The organs from a single animal representing each group were given for histological studies. PAS staining was done to estimate the glycogen content in liver specimens. Also, MTS staining was performed in liver tissues to estimate fibrosis. The specimens were submitted to Dr Vamshi’s Biological Sciences and Research Center, Rajajinagar, Bangalore for detailed examination of the tissues.

### Statistical analysis

All results are expressed as the means ± standard deviations (SDs). A one-way ANOVA test was performed to determine the significance of test samples compared to the controls and a value of p<0.05 was considered as significant.

## Results

### *Lodhrasavam* protects against palmitic acid–induced cytotoxicity and hepatic steatosis by reducing lipid accumulation and downregulating lipogenic gene expression in HepG2 cells

The cytoprotective and anti-steatotic effects of Lodhrasavam (LS) were evaluated using a palmitic acid (PA)-induced in vitro model of MASLD in HepG2 cells. Preliminary cytotoxicity testing revealed that LS is largely non-toxic to HepG2 cells up to 64μg GAE/ml, with only the highest dose (128μg GAE/ml) showing a reduction in cell viability to 25.67% ± 10.48%. Lower concentrations (2μg GAE/ml) maintained high cell viability (92.66% ± 9.43%) (Fig.1B).

**Figure 1:**
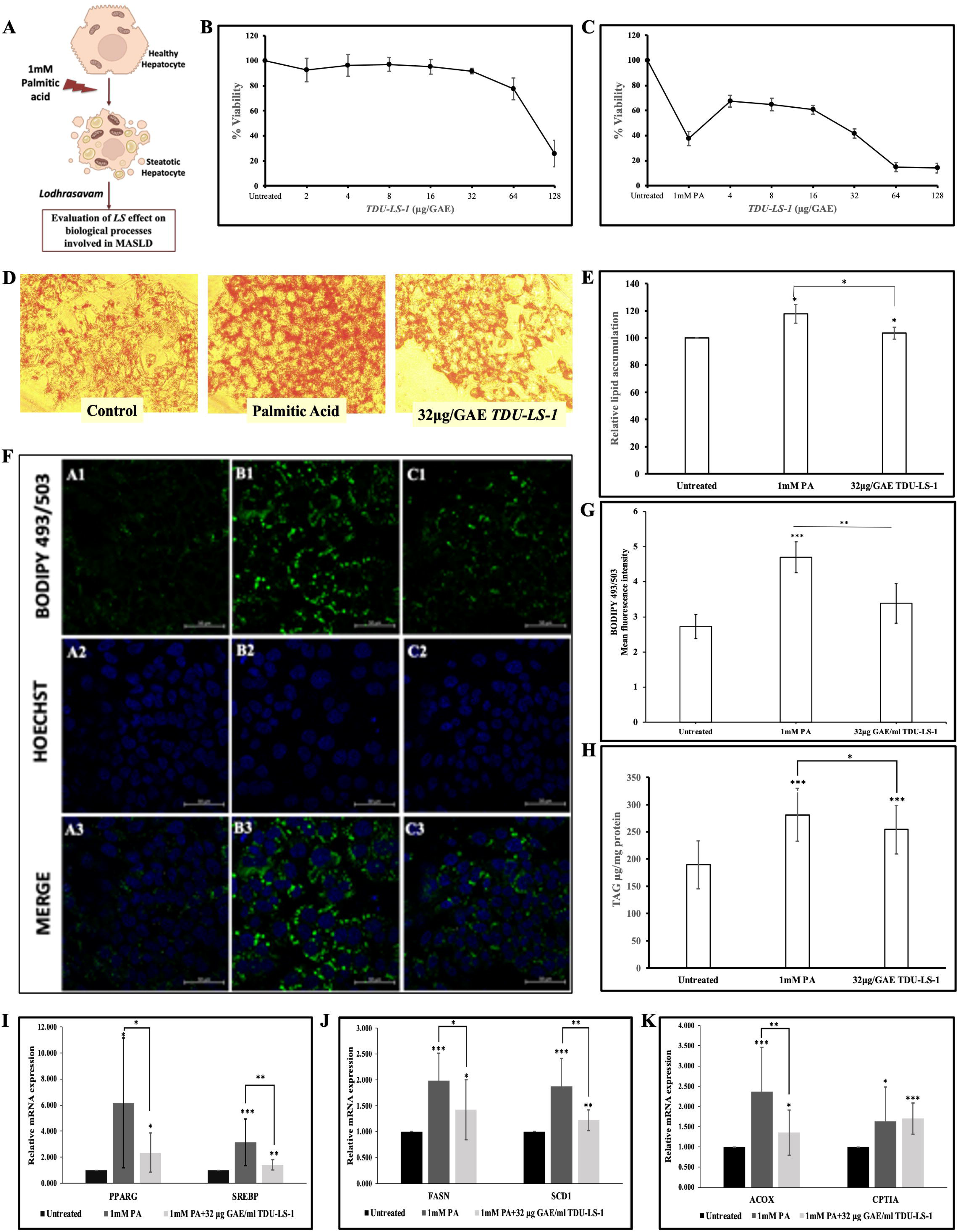
Lodhrasavam (LS) attenuates palmitic acid (PA)-induced cytotoxicity, lipid accumulation, and lipogenic gene expression in HepG2 cells. (A) Schematic representation of the HepG2 cell steatotic model; (B) Cell viability assessed by MTT assay following treatment with various concentrations of LS (2–128μg GAE/ml) for 24 h, indicating dose-dependent cytocompatibility; 128μg GAE/ml showed significant cytotoxicity; (C) Viability of HepG2 cells exposed to 1 mM PA for 24 h with or without LS post-treatment. LS (4–32μg GAE/ml) significantly reversed PA-induced cytotoxicity; (D-E) Oil Red O staining quantifying intracellular lipid content, with LS treatment reducing PA-induced lipid accumulation; (F-G) BODIPY 493/503 staining showing increased lipid droplet accumulation in PA-treated cells (B1-3) compared to control cells (A1-3), which was reduced by 32μg GAE/ml LS (C1-3); (H) Triglyceride (TG) assay showing a 1.10-fold reduction in TG content upon LS treatment versus PA alone. (I-K) qRT-PCR analysis showing PA-induced upregulation of PPARγ, SREBP-1c, FASN, SCD1 and ACOX was significantly downregulated and CPT1*α* levels increased following LS treatment at 32μg GAE/ml. Data are presented as mean ± SD from three independent experiments. *p < 0.05, **p < 0.005, ***p < 0.001 versus PA-treated group.

Exposure of HepG2 cells to 1 mM PA for 24 h significantly reduced cell viability to 37.64% ± 5.71%, indicating cytotoxicity along with lipid and triglyceride accumulation. Post-PA treatment with LS demonstrated dose-dependent cytoprotection. Specifically, LS at concentrations of 4, 8, 16, and 32μg GAE/ml significantly increased cell viability to 67.46% ± 4.9%, 64.81% ± 5.06%, 60.69% ± 3.44%, and 41.64% ± 3.70%, respectively, reversing PA-induced cytotoxic effects. Although 64μg GAE/ml LS was non-toxic in isolation, it was not effective in reversing PA-induced cytotoxicity (Fig. 1C).

To assess the impact of LS on hepatic lipid accumulation, Oil Red O (ORO) staining corroborated this, with a 1.14-fold reduction in lipid droplet accumulation in LS-treated cells compared to PA-treated cells (Fig 1D-E). BODIPY 493/503 staining was performed. PA-treated cells exhibited a 1.72-fold increase in mean fluorescence intensity, indicating increased lipid droplet accumulation. Treatment with 32μg GAE/ml LS reduced fluorescence intensity by 0.72-fold, indicating lipid-lowering activity (Fig. 1F-G). Further, PA-induced triglyceride (TG) accumulation was reduced by 1.10-fold in the LS-treated group, indicating an improvement in intracellular lipid homeostasis (Fig. 1H).

At the molecular level, LS treatment modulated the expression of key lipogenic genes implicated in hepatic steatosis. PA exposure significantly upregulated PPARγ (6.2-fold), SREBP-1C (3.2-fold), FASN (2.0-fold), SCD1 (1.9-fold), and ACOX1 (2.36-fold). Treatment with 32μg GAE/ml LS significantly attenuated these elevations, reducing PPARγ by 2.62-fold, SREBP-1C by 2.21-fold, FASN by 1.39-fold, SCD1 by 1.54-fold, and ACOX1 by 1.75-fold, compared to PA-treated cells (Fig. 1I-K).

### *Lodhrasavam* exhibits potent antioxidant activity and reduces oxidative stress and inflammatory markers

The antioxidant potential of Lodhrasavam (LS) was evaluated using a DPPH radical scavenging assay. The results demonstrated a concentration-dependent increase in free radical scavenging activity, with the highest tested concentration (128μg GAE/ml) showing a significant 90.40% ± 1.39% inhibition of DPPH radicals. In contrast, the lowest concentration (2μg GAE/ml) exhibited 15.33% ± 5.38% inhibition (Fig. 2A). The IC₅₀ value of LS was determined to be 16.45μg GAE/ml, which was lower than that of ascorbic acid (24.51μg/ml), indicating higher antioxidant potency. Furthermore, the Ascorbic Acid Equivalent Antioxidant Capacity (AEAC) of LS was calculated to be 1.49, underscoring its strong radical-neutralizing capability.

**Figure 2:**
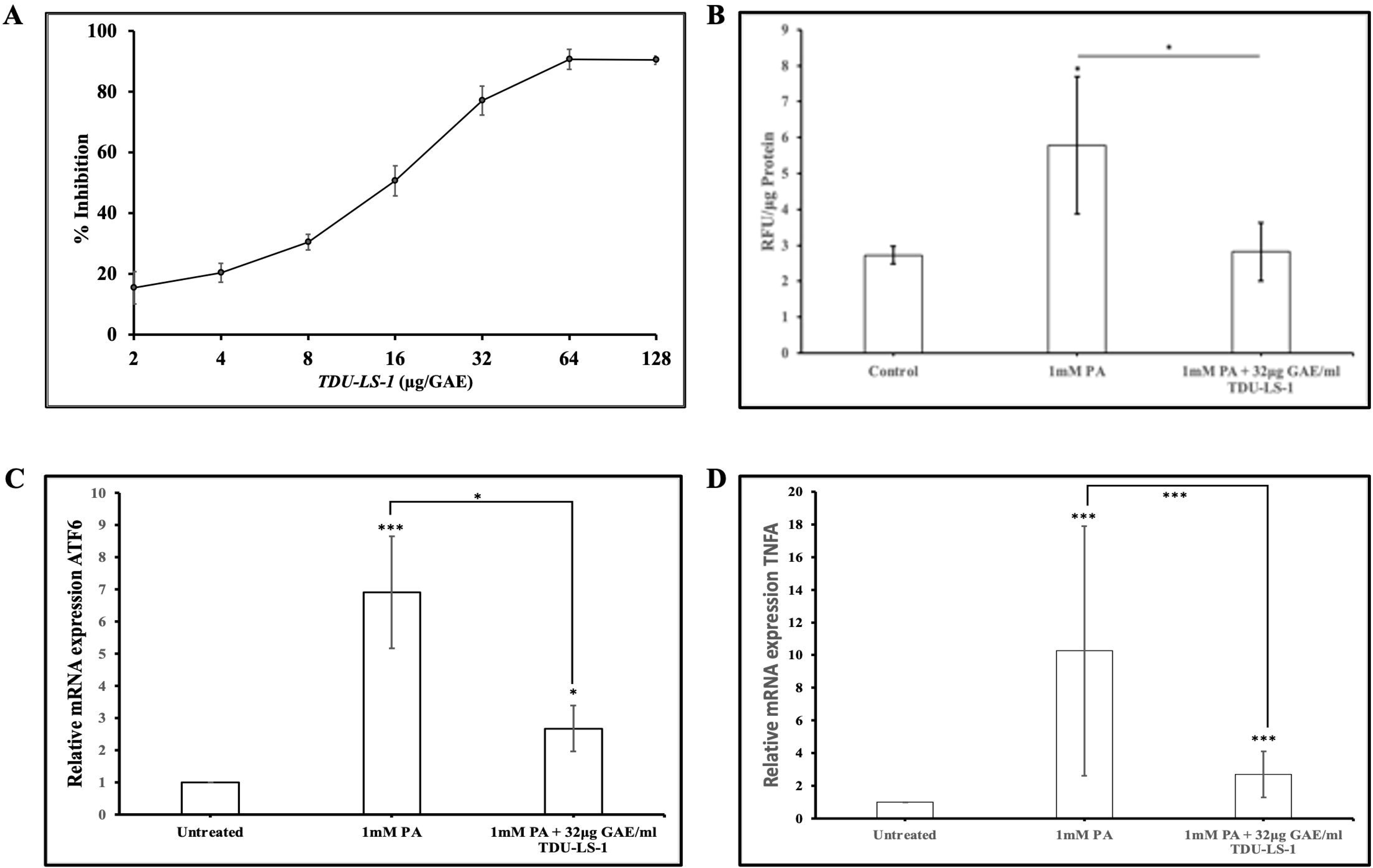
*LS exhibits potent antioxidant activity and attenuates PA-induced oxidative stress and inflammation.* (A) DPPH radical scavenging activity of LS at various concentrations (2–128µg GAE/ml), showing a dose-dependent increase in free radical neutralization. LS exhibited an IC₅₀ of 16.45µg GAE/ml, lower than that of ascorbic acid (24.51µg/ml), indicating higher antioxidant potency; (B) Intracellular ROS levels in HepG2 cells exposed to 1 mM palmitic acid (PA) with or without LS co-treatment (32µg GAE/ml). LS significantly reduced PA-induced ROS accumulation (∼2-fold decrease); Relative gene expression of inflammatory markers TNF-α (D) and ATF6 (C) following PA exposure and LS treatment, showing significant downregulation with LS, indicative of anti-inflammatory effects. Data are expressed as mean ± SD from at least three independent experiments. *p < 0.05, **p < 0.005, ***p < 0.001, versus PA-treated group.

**Figure 3:**
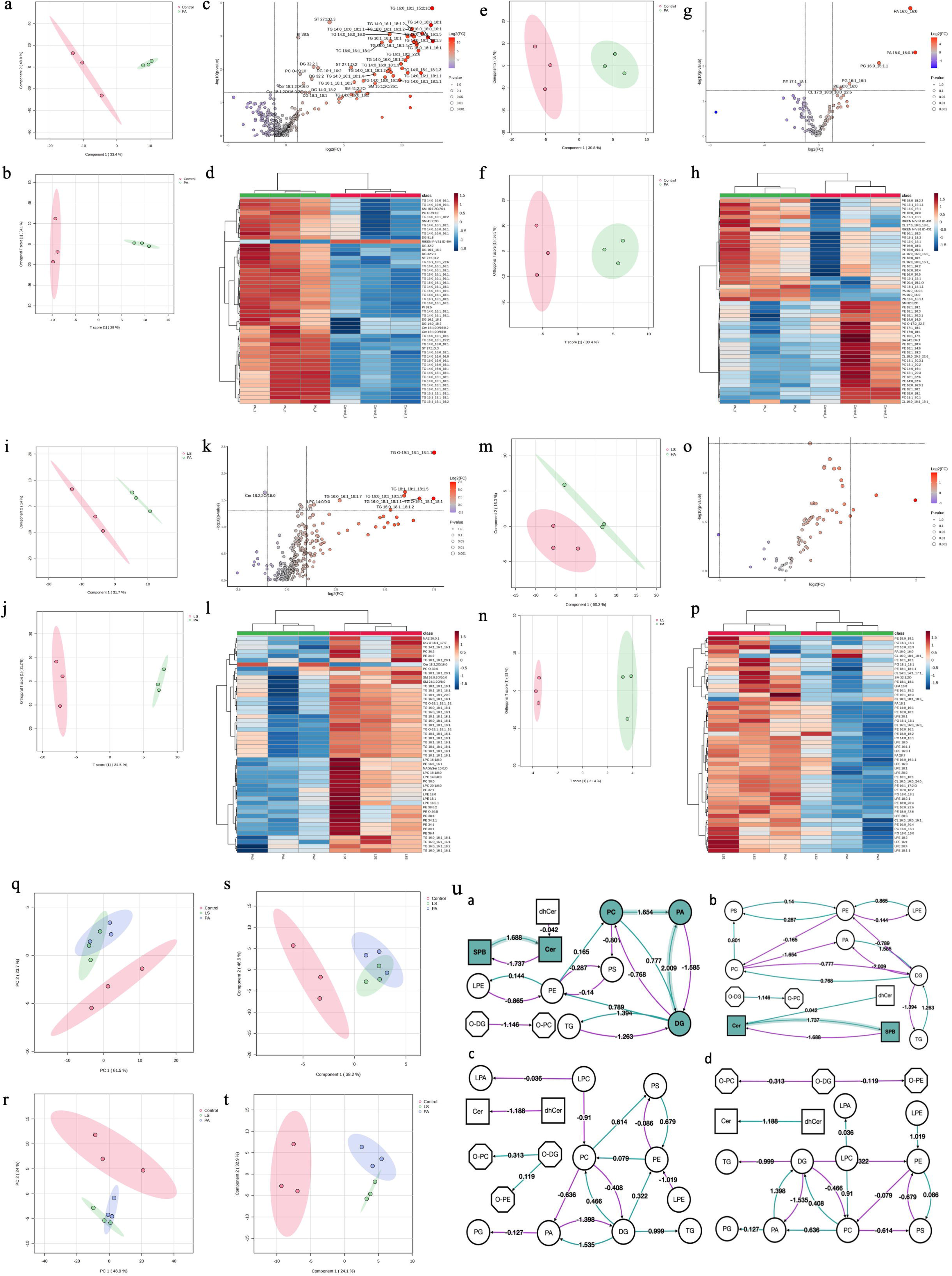
*LS Modulates Palmitate-Induced Lipidomic Alterations.* (a, e) PLS-DA score plots showing clear separation between Control and PA groups in positive (a) and negative (e) ion modes, indicating palmitate-induced lipidomic shifts. (b, f) OPLS-DA plots further refining group separation between Control and PA in positive (b) and negative (f) modes. (c, g) Volcano plots and (d, h) heat maps showing significantly upregulated TGs, sterols, and phospholipids in PA vs Control in positive (c, d) and negative (g, h) modes. (i, m) PLS-DA plots showing moderate separation between PA and LS groups in positive (i) and negative (m) ion modes, reflecting partial restoration of lipid profiles by Lodhrasavam. (j, n) OPLS-DA plots for PA vs LS comparison in positive (j) and negative (n) modes. (k, o) Volcano plots and (l, p) heat maps showing downregulation of TGs and ceramides in LS-treated cells compared to PA in positive (k, l) and negative (o, p) ion modes. (q–t) PCA and PLS-DA plots comparing global lipidomic variance across Control, PA, and LS groups; no statistically significant global separation observed. (u) BioPAN pathway analysis showing lipid metabolism shifts in PA vs Control and their normalization by Lodhrasavam. (ua–ud) Specific BioPAN modules depicting active lipid class conversions: (ua) PA-induced lipogenesis and ceramide accumulation, (ub) suppressed phospholipid and sphingolipid turnover, (uc) LS-induced recovery of TG and PC biosynthesis, (ud) normalization of membrane lipid remodeling.

In cellular assays, treatment with 1 mM palmitic acid (PA) significantly elevated intracellular reactive oxygen species (ROS) levels by 2.12-fold compared to untreated controls. However, co-treatment with LS at 32μg GAE/ml reduced ROS levels by approximately 2-fold compared to PA-only treated cells, highlighting the ability of LS to mitigate oxidative stress (Fig 2B). Additionally, LS significantly downregulated the expression of pro-inflammatory markers TNF-α and ATF6, suggesting a strong anti-inflammatory effect (Fig 2C-D).

### *Lodhrasavam* Attenuates Palmitate-Induced Lipotoxicity Through Distinct Modulation of Lipidomic Signatures Revealed by Multivariate and Univariate Analyses

Multivariate modelling revealed distinct lipidomic shifts following PA and LS treatments. PLS-DA revealed clear separation between Control and PA groups in both ionization modes, reflecting robust palmitate-induced lipidomic remodeling. In positive mode, Components 1 and 2 explained 82.25% of the variance (33.45% and 48.8%, respectively) (3a), while in negative mode, they accounted for 86.8% (30.8%, 56.0%) (3e). Top VIP-scoring lipids in positive mode included saturated and monounsaturated triacylglycerols (TGs) such as TG 16:0_18:1_15:2;1O and TG 14:0_16:0_16:0 (VIP > 1.85), along with non-TG lipids (DG 32:2, PI 38:5, ST 27:1;O.3), indicating enhanced lipogenesis and disrupted sterol/phosphoinositide metabolism. In negative mode, VIP lipids were dominated by phospholipids including PA 16:0_16:0, PG 16:0_16:1.1, and PE 17:1_18:1, reflecting altered membrane dynamics and mitochondrial stress. OPLS-DA further refined group separation, attributing 28.0% (positive) and 30.4% (negative) of variance to treatment, with 54.1% and 56.5% orthogonal variation (3b and 3f). Volcano plot analysis supported these findings, identifying TGs (log₂FC > 12, *p* < 0.001), sterols, and DGs as highly elevated in PA vs Control (positive mode), and strong upregulation of PA 16:0_16:0 and PG 16:0_16:1.1 in negative mode.

Lodhrasavam reverses palmitate-induced lipidomic perturbations as observed in PLS-DA analysis of PA vs LS groups demonstrated moderate but clear separation in both ion modes, with Components 1 and 2 explaining 31.7% and 14.0% (positive, 3i) and 60.2% and 16.3% (negative, 3m) of total variance. Top VIP-scoring features in positive mode included TG O-19:1_18:1_18:1.1, TG 18:1_18:1_18:1.5, and LPCs (14:0/0:0, 18:1/0:0), suggesting modulation of neutral and lysolipid pools. Cer 18:2;2O/16:0 was downregulated (VIP = 1.75, *p* = 0.023), highlighting anti-lipotoxic effects. In negative mode, high-ranking lipids (VIP > 1.5) included PG 16:1_16:1, PE 16:0_16:1, and CL 16:0_16:0_16:0_18:0, indicating restoration of phospholipid and mitochondrial lipid profiles. OPLS-DA showed 24.5% (positive, 3j) and 21.4% (negative, 3n) of variance linked to treatment, with orthogonal components capturing 21.2% and 53.0%, respectively. These results highlight the ability of LS to modulate both storage and structural lipids, particularly TGs, ceramides, phospholipids, and lysolipids.

Univariate analysis confirms modulation of key lipid species. Volcano plot analysis of Control vs PA (positive mode, 3c) revealed significant enrichment of triacylglycerols (TGs) and sterol lipids, with several TGs (e.g., TG 16:0_18:1_15:2;1O, TG 14:0_16:0_18:1) showing extremely high fold changes (log₂FC > 12, *p* < 0.001), and sterol derivatives (e.g., ST 27:1;O.3) markedly upregulated, indicating enhanced lipogenesis and altered sterol metabolism. Elevated diacylglycerols and sphingolipids (e.g., DG 32:2.1, Cer 18:1;2O/16:0) suggested membrane stress and lipotoxic signaling. In negative mode (3g), phosphatidic acid (PA 16:0_16:0, FC = 107.2, *p* = 0.0002) and PG 16:0_16:1.1 (FC = 21.6, *p* = 0.0081) were significantly upregulated, while PE 17:1_18:1 was downregulated (FC = 0.37, *p* = 0.034), indicating disrupted phospholipid metabolism and mitochondrial stress—findings corroborated by OPLS-DA VIP profiles. Volcano plot analysis of PA vs LS (positive mode, 3k) identified significantly downregulated TGs, including TG O-19:1_18:1_18:1.1 (log₂FC = 7.51), TG 16:0_18:1_18:1.1 (log₂FC = 6.76), and TG 18:1_18:1_18:1.5 (log₂FC = 6.03), consistent with reduced lipid accumulation. Cer 18:2;2O/16:0 was also significantly downregulated (log₂FC = –1.12, *p* = 0.023). In contrast, negative mode analysis did not yield statistically significant results, suggesting a more pronounced intervention effect on neutral lipids (3o).

Global lipidome variation is moderate and not statistically significant at the multivariate level. Pairwise PERMANOVA comparisons between groups (Control vs PA, Control vs LS, PA vs LS) showed moderate separation (R² = 0.31–0.39) in positive mode, but none reached statistical significance after FDR correction (*p.adj* ≥ 0.30). This suggests that while specific lipid species were differentially regulated, the overall global lipidomic profile remained statistically similar across treatments (3q-t). Nevertheless, supervised multivariate models like PLS-DA and OPLS-DA effectively captured treatment-driven lipidomic shifts.

BioPAN analysis revealed significant lipid remodeling in PA-treated cells and its modulation by Lodhrasavam (3u). In the PA vs Control comparison, the most active lipid subclass conversion was DG→PA (Z = 2.009; *DGK family*), along with enhanced SPB→Cer (Z = 1.688; *CERS1–6*) and PC→PA (Z = 1.654; *PLD1/2*), indicating elevated TAG precursor formation and ceramide accumulation (3ua). Lipid species-level reactions such as PC(32:2)→DG(32:2)→TG(50:2) and FA(16:0)→FA(16:1) (Z = 2.326; *SCD3*) further support increased lipogenesis and desaturation. Conversely, PA suppressed phospholipid re-synthesis pathways, notably DG→PC and PE→PC conversions (Z > 2.3), involving *CHPT1*, *PEMT*, and *CEPT1*. The Cer→SPB reaction (Z = 1.737; *ASAH1–2*, *ACERs*) was also inhibited, suggesting impaired sphingolipid turnover (3ub). Compared to PA, Lodhrasavam (LS) treatment enhanced DG(32:2)→TG(50:3) (Z = 2.4; *DGAT2*) and PE(38:4)→PC(38:4) (Z = 1.706; *PEMT*), reflecting restored glycerolipid and phospholipid metabolism (3uc). Simultaneously, LS suppressed excessive PE/LPC→PC conversions (Z = 2.729), indicating normalization of membrane lipid remodeling via *PEMT* and *LPCATs* (3ud). These findings highlight that Lodhrasavam mitigates PA-induced lipid dysregulation by curbing lipotoxic TAG synthesis and restoring phospholipid homeostasis.

### *Lodhrasavam* ameliorates HFD-induced hepatic steatosis, and dyslipidemia through modulation of metabolic parameters in rats

Over a 90-day HFD induction period followed by a 30-day treatment phase, body weight was monitored to assess obesity progression and therapeutic impact. Rats in the HFD group exhibited significant weight gain, while those treated with either LS or PIOZ demonstrated a marked and statistically significant reduction in body weight during the treatment period (p < 0.05; p < 0.005) (Fig. 4C-D). These results indicate that LS mitigates diet-induced weight gain and may exert anti-obesity effects.

**Figure 4:**
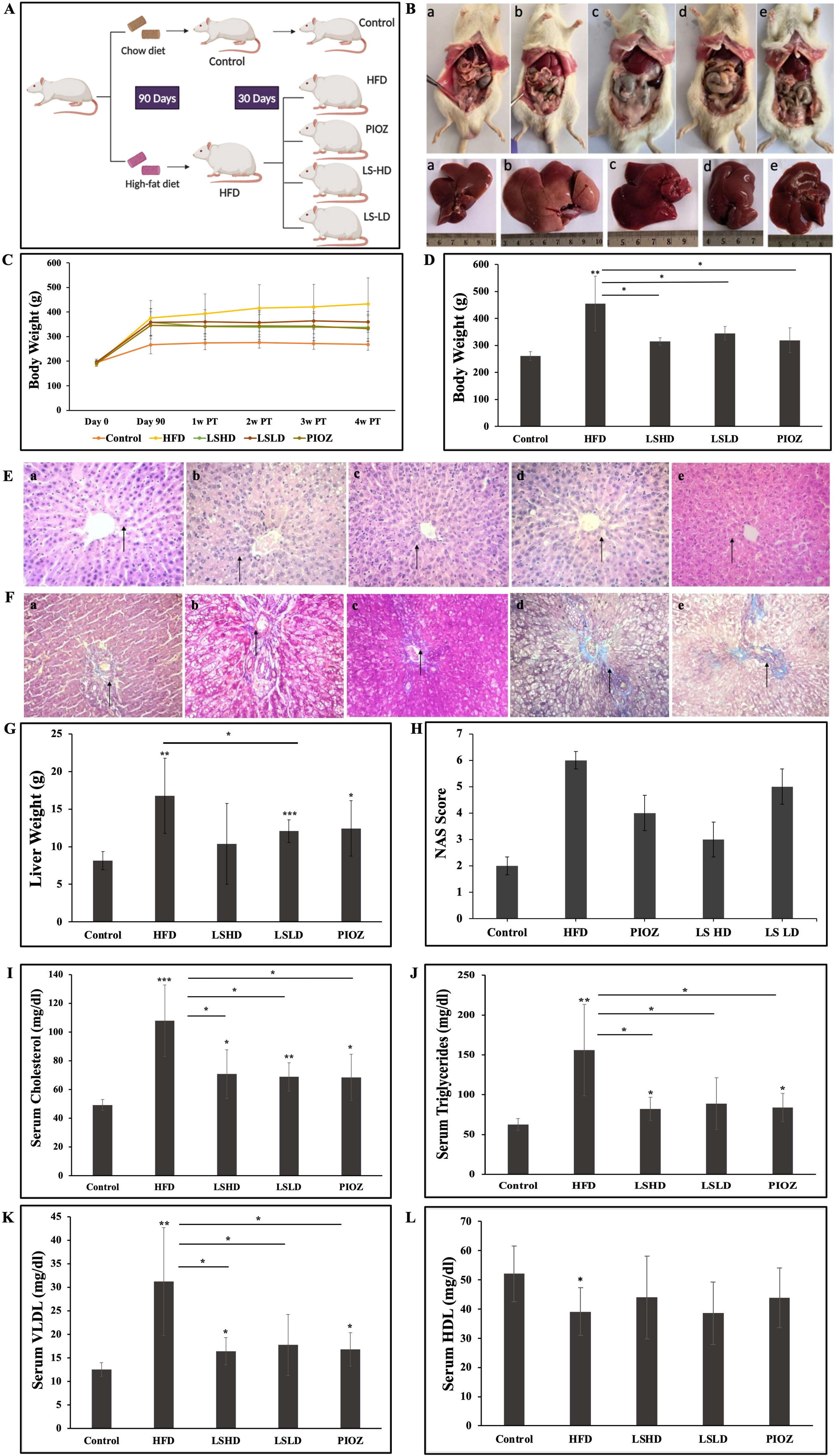
*LS mitigates HFD-induced hepatic steatosis, and dyslipidemia in rats.* (A) Schematic representation of *in vivo* model of MASLD; (B) Images of dissected rat with respective livers (a) control, (b) HFD, (c) LSHD, (d) LSLD and (e) PIOZ; (C-D)Body weight changes during the 30-day treatment period following 90 days of HFD induction. LS-treated (LSLD, LSHD) and PIOZ-treated groups showed significant reductions in weight gain compared to the HFD group; (E-F) Representative H&E-stained and MTS stained liver sections depicting histological improvements in steatosis, lobular inflammation, and hepatocyte ballooning in LS and PIOZ groups compared to HFD; (G) Liver weight measurements showing reduced hepatic hypertrophy in all treatment groups, with LSLD exhibiting the most pronounced effect; (H) NAFLD Activity Score (NAS) indicating significant amelioration of hepatic damage across all treatment groups; (I-K) Serum lipid parameters showing significantly reduced total cholesterol, triglycerides, and VLDL levels in LS and PIOZ groups compared to HFD; (L) HDL levels showed a modest, non-significant increase. Data are presented as mean ± SD (n = 6 per group); *p < 0.05, **p < 0.01, ***p < 0.001.

Liver weight, a surrogate marker for steatosis and hepatomegaly, was significantly reduced in all treatment groups (LSHD, LSLD, and PIOZ) compared to the HFD group. Among them, the LSLD group exhibited the most pronounced decrease in liver weight, indicating its potential efficacy in reversing hepatic lipid accumulation and associated pathological enlargement (Fig. 4G).

Histological examination of liver tissue using hematoxylin and eosin (H&E) staining and MTS staining revealed substantial reductions in macrovesicular steatosis, lobular inflammation, and hepatocyte ballooning in the LSHD, LSLD, and PIOZ-treated groups compared to the HFD group (Fig. 4E-F). These improvements were quantified using the NAFLD Activity Score (NAS), which showed a statistically significant reduction in all treatment groups, indicating amelioration of diet-induced hepatic damage (Fig. 4H). The LSHD and LSLD treatments particularly demonstrated strong hepatoprotective effects.

Serum biochemical analysis revealed that LS treatment significantly improved the lipid profile disrupted by HFD feeding. Total cholesterol and triglyceride levels were significantly reduced in treated groups compared to the HFD group, reflecting effective attenuation of hyperlipidaemia (Fig. 4-1-J). VLDL levels, a marker of hepatic triglyceride output and atherogenicity, were also significantly decreased (Fig. 4K). Although HDL levels showed a modest upward trend post-treatment, the increase was not statistically significant (Fig. 4L). These results suggest that LS improves lipid metabolism, primarily through the reduction of cholesterol, triglycerides, and VLDL.

### *Lodhrasavam* attenuates adipogenesis and lipid metabolism by inhibiting lipogenesis, lipase activity, and reducing adipose tissue mass

LS demonstrated robust anti-adipogenic effects in vitro and in vivo by targeting key processes involved in lipid accumulation and metabolism. In 3T3-L1 preadipocytes induced with MDI, LS significantly inhibited adipogenesis, as evidenced by a marked reduction in both intracellular triglyceride (TAG) levels and lipid droplet accumulation. MDI treatment alone caused a 7.5-fold increase in TAG levels relative to the control, whereas treatment with 32µg GAE/ml LS led to a 2.8-fold decrease in TAG content compared to the MDI group. Similarly, LS treatment resulted in a 2.12-fold reduction in lipid droplet accumulation, counteracting the 2.13-fold increase observed in MDI-treated cells (Fig. 5D-F).

**Figure 5:**
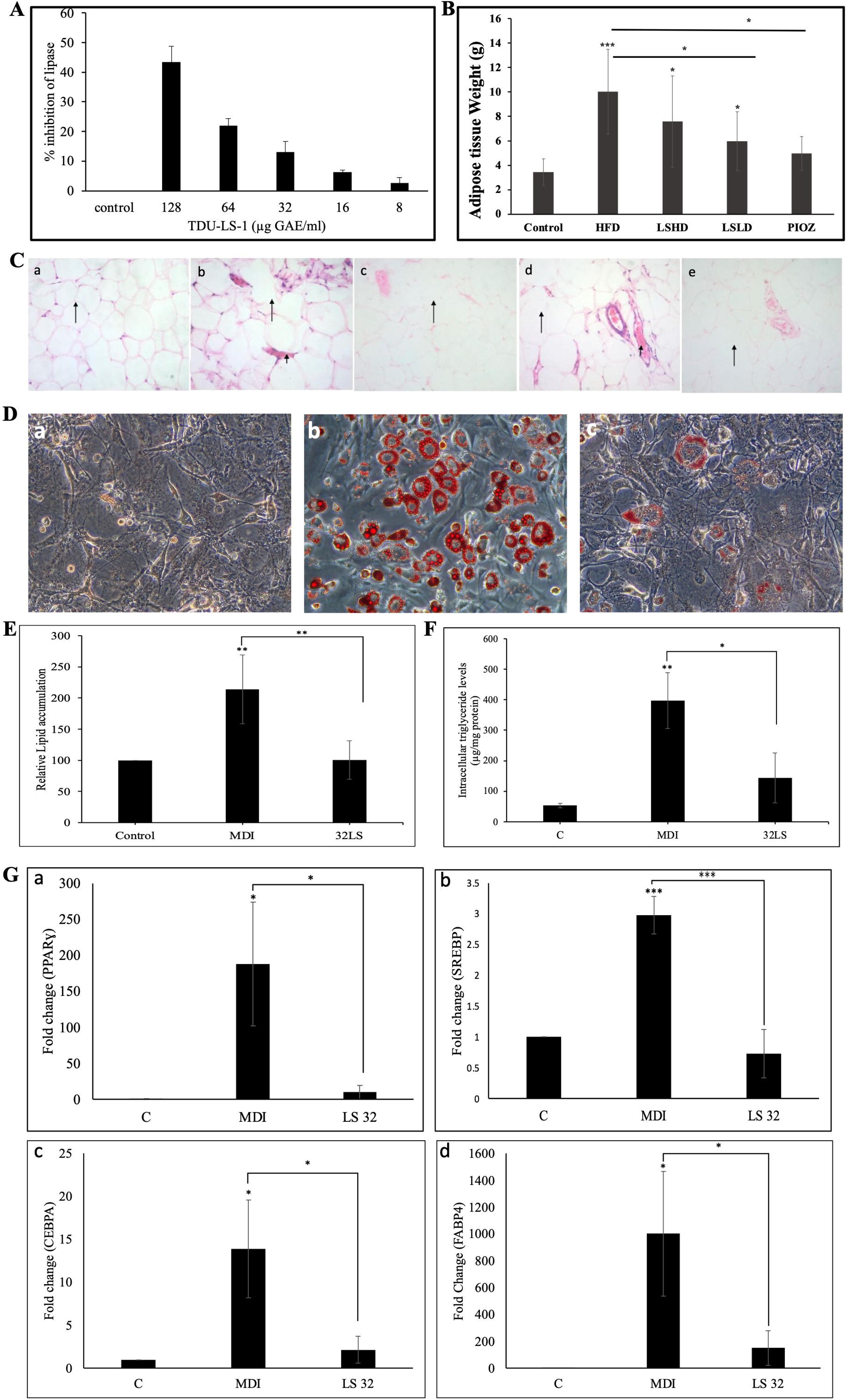
LS attenuates adipogenesis, lipogenesis, and adipose tissue hypertrophy through in vitro and in vivo modulation of lipid metabolism. (A) Pancreatic lipase inhibition assay results indicating dose-dependent inhibition by LS, with a maximum of 43 ± 5% inhibition at 128µg GAE/ml; (B) Quantification of adipose tissue mass indicating significant reduction in all treatment groups, most notably LSLD and PIOZ; (C) Representative H&E-stained adipose tissue sections from in vivo models showing preservation of adipocyte architecture in LS- and PIOZ-treated groups compared to HFD; (D-E) Lipid droplet accumulation measured by Oil Red O staining (a-control, b-MDI, c-LS), with LS significantly reversing MDI-induced lipid accumulation; (F) Quantification of intracellular triglyceride (TAG) levels in 3T3-L1 preadipocytes showing LS-induced reduction in MDI-stimulated adipogenesis; (G a-d) Relative mRNA expression of adipogenic and lipogenic genes (*Pparγ, Srebp, Cebpα, Fabp4*) demonstrating marked downregulation in LS-treated cells. Data are expressed as mean ± SD (n = 6 for in vivo); *p < 0.05, **p < 0.01, ***p < 0.001.

Gene expression analysis further elucidated LS’s mechanistic role in inhibiting adipogenesis. LS significantly downregulated the transcription of adipogenic master regulators and lipogenic genes. Compared to MDI-only treatment, LS reduced the mRNA expression levels of *Pparγ* by 18.9-fold, *Srebp* by 4.25-fold, *Cebpα* by 6.5-fold, and *Fabp4* by 6.7-fold was observed (Fig. 5G), suggesting LS may modulate lipid metabolism pathway.

Histopathological examination of adipose tissue from in vivo models supported these in vitro findings. Haematoxylin and eosin (H&E) staining revealed well-preserved adipocyte architecture in both LSHD and PIOZ-treated groups, with no signs of vascular congestion or inflammatory infiltration, in stark contrast to the HFD group (Fig. 5C). Furthermore, quantitative assessment of adipose tissue mass showed a significant reduction across all treatment groups compared to HFD, with the LSLD and PIOZ groups exhibiting the most prominent effects (Fig. 5B). These findings underscore the capacity of LS to prevent diet-induced adipose tissue expansion and associated histological damage.

In addition to its anti-adipogenic effects, LS exhibited dose-dependent inhibition of pancreatic lipase, a key enzyme responsible for the hydrolysis of dietary triglycerides into absorbable fatty acids. At its highest tested concentration (128µg GAE/ml), LS inhibited pancreatic lipase activity by 43 ± 5% (Fig. 5A), suggesting its potential to reduce dietary fat absorption and thereby combat hyperlipidaemia and obesity-related complications.

### *Lodhrasavam* ameliorates glucose dysregulation and pancreatic dysfunction in HFD-induced MASLD potentially via enhanced insulin and GLP-1 secretion

To assess the efficacy of Lodhrasavam in modulating glucose metabolism under high-fat diet (HFD)-induced metabolic dysfunction, an Oral Glucose Tolerance Test (OGTT) was conducted. Blood glucose levels were measured at 0, 15, 30, 60, and 120 minutes post-glucose administration. The treatment groups exhibited a marked improvement in glucose clearance compared to the HFD group, as reflected by significantly lower blood glucose levels between 30 and 120 minutes (Fig. 6C). Quantitative assessment of glucose tolerance, based on the area under the curve (AUC) of the OGTT profile, confirmed these findings (Fig. 6D). Notably, animals treated with both high-dose (LSHD) and low-dose (LSLD) Lodhrasavam formulations showed significantly enhanced glucose disposal, suggesting improved glucose homeostasis, likely attributable to increased peripheral glucose utilization and improved insulin dynamics.

**Figure 6:**
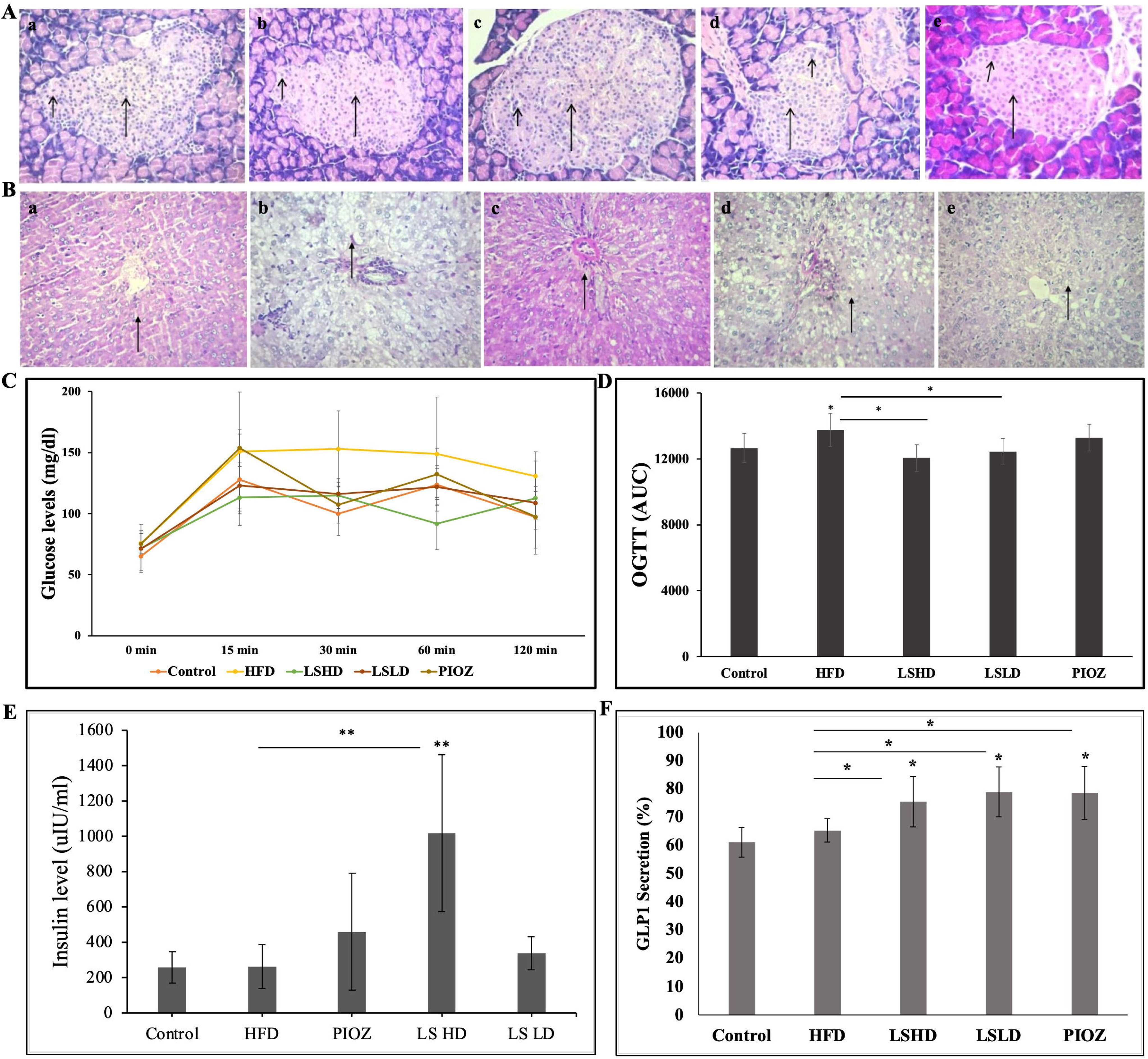
*LS improves glucose homeostasis and pancreatic function in HFD-induced MASLD via enhanced insulin and GLP-1 secretion.* (A) Representative H&E-stained pancreatic sections showing restoration of islet architecture and β-cell mass in LS-treated rats compared to HFD; (B) PAS staining of liver samples showing increased glycogen content LSHD when compared to HFD; (C) Oral Glucose Tolerance Test (OGTT) showing blood glucose levels at 0, 15, 30, 60, and 120 minutes post-glucose administration; (D) Area Under the Curve (AUC) analysis of OGTT indicating significantly improved glucose clearance in LSHD and LSLD groups vs HFD; (E) Serum insulin levels showing a significant increase, especially in the LSHD group; (F) Serum GLP-1 levels significantly elevated in all treatment groups, with the LSHD group demonstrating the highest increase. All data are presented as mean ± SD (n = 6); . *p < 0.05, **p < 0.01.

Histological analysis of the pancreas using haematoxylin and eosin (H&E) staining provided further insight into pancreatic integrity and function (Fig. 6A). In the HFD group, pancreatic tissue revealed a reduction in islet number and β-cell mass, with β-cells constituting approximately 55% of the islet population. In contrast, the LSHD group exhibited a substantial restoration of islet architecture and β-cell mass, with β-cells comprising about 65% of the islet population—indicating a regenerative or protective influence of the formulation. While the LSLD and PIOZ-treated groups showed slight morphological improvements, the extent of β-cell restoration was less pronounced.

Functional assessment corroborated the histological data, as all treated groups demonstrated increased insulin secretion relative to the HFD control, with the LSHD group showing a statistically significant rise (Fig. 6E). This enhancement in insulin output strongly correlated with the observed increase in β-cell mass, supporting the hypothesis that Lodhrasavam may exert trophic effects on pancreatic β-cells under metabolic stress.

Additionally, GLP-1 secretion, a key modulator of glucose-dependent insulin release and β-cell preservation was significantly upregulated in all treatment groups compared to the control and HFD groups (Fig. 6F). The LSHD group demonstrated the highest elevation in GLP-1 levels, further reinforcing the formulation’s role in enhancing incretin signalling, which is central to maintaining glycaemic control and slowing MASLD progression.

## Discussion

Metabolic dysfunction-associated steatotic liver disease (MASLD), previously known as NAFLD, is a complex, multifactorial condition closely linked to obesity, insulin resistance, oxidative stress, and chronic inflammation. Current single molecule therapeutic approaches have shown limited efficacy in addressing this multifaceted disease, prompting interest in polyherbal *Ayurveda* formulations with multitargeted mechanisms. In previous studies conducted in our lab *Lodhrasavam* has shown potential anti-diabetic and anti-adipogenic effects. It inhibited α-amylase and α-glucosidase activity by 90% and 78% respectively and reduced adipocyte differentiation significantly (Butala et al., 2017). It has not been explored for its anti-steatosis and antioxidant activities. In this study, we comprehensively evaluated the therapeutic potential of *Lodhrasavam* (LS), a classical *Ayurveda* formulation, in both in vitro and in vivo models of MASLD.

Several *Ayurveda* formulations have been studied for their effect on improving liver health and overall systemic metabolism. In a case study conducted by Sahu et al., 2022, it was found that *Ayurveda* formulations (*Avipattikara Churna, Punarnava Mandura, Sankha Bhasma, Tarunikusumakara Churna, Triphala Churna* and *Kutaki Churna*) have immense potential in treating grade 2 fatty liver disease. A similar effect was seen in a randomized placebo-controlled clinical trial of a multiherbal formulation, *Sharapunkhadi* powder on subjects with grade 1-3 fatty liver disease (Remya et al., 2020). Livogrit, a tri-herbal *Ayurvedic* medicine, was studied for its hepatoprotective effect on methionine and cysteine deficient NASH models in both HepG2 spheroids and rat primary hepatocytes. It reduced fat accumulation and ROS production and increased lipolysis hence attenuating NASH (Balkrishna et al., 2022).

MASLD is characterized by the accumulation of lipid droplets (LDs) in the hepatocytes, referred to as hepatic steatosis. It is considered as the “first-hit” in the development of MASLD. LDs can accumulate in the liver via three mechanisms: a) flux of free fatty acids (FFAs) from adipose tissue due to increased lipolysis, increased *de novo lipogenesis* (DNL), and dietary fat intake; b) decreased export of triglycerides and c) reduced beta-oxidation. Elevated plasma free fatty acid (FFA) levels play an etiological role in the pathogenesis of MASLD. In particular, the saturated fatty acid palmitate, which makes up 30%-40% of high plasma FFA concentration, is a major contributor to an increase in intrahepatic triglyceride (Cacicedo et al., 2005; Schroeder-Gloeckler et al., 2007).

In the present study, 1 mM PA was used to induce cellular steatosis (Gómez-Lechón et al., 2007). *In vitro* investigations using HepG2 hepatocytes demonstrated that LS protects against palmitic acid (PA)-induced cytotoxicity and steatosis. LS significantly restored cell viability in a dose-dependent manner and reduced intracellular lipid droplet and triglyceride accumulation, indicating potent anti-steatotic activity. To investigate the mechanism of LS inhibition of hepatic steatosis, we evaluated the expression of key genes involved in lipogenesis, including SREBP-1c, FASN and SCD-1 (Parlati et al., 2021). SREBP-1c is a master regulator of lipogenesis, known to transcriptionally regulate downstream genes involved in de novo lipogenesis, including those encoding ACC and FAS (Osborne, 2000). In a study by Motoyuki et al., 2007, it showed an increase in expression of SREBP-1c, ACC and FAS in MASLD patients. It has been reported that SREBP-1c activation is associated with increased lipogenesis in MASLD (Yan et al., 2014). In mice, HFD caused fat accumulation and induced SREBP-1c mRNA expression which further contributes to the progression of hepatic steatosis (Kim et al., 2010; Zhu et al., 2019). Li et al., 2011 showed that inhibiting SREBP-1c activity can exert an anti-hepatic steatosis effect in diet-induced obese mice. SCD-1 is an endoplasmic reticulum bound rate limiting enzyme that catalyzes the formation of mono-unsaturated fatty acids from saturated fatty acids (Paton and Ntambi, 2009) Liver specific knock-out of SCDI in mice fed with high-carbohydrate diet showed reduction in hepatic steatosis (Miyazaki et al., 2007) (Jiang et al., 2005) in their experiments using antisense oligonucleotide inhibitor of SCD1 showed inhibition of lipogenesis and promoted lipid oxidation in both primary mouse hepatocytes and HFD fed mice. In the current study, 24hr incubation with 1mM PA increased the expression of these genes. Following which LS significantly downregulated the expression of SREBP-1c, FAS and SCD-1.

The hepatic PPARγ plays a putative role in the progression of fatty liver disease in the MASLD patients. PPAR-γ is up-regulated in the liver of obese patients with MASLD, and recently, the expression of PPAR-γ is considered as an additional reinforcing lipogenic signal, assisting SREBP-1c to trigger the development of hepatic steatosis (Pettinelli and Videla, 2011). We showed that PA increased PPAR-γ mRNA expression in HepG2 cells, which was significantly reduced by LS.

When there is an excess of fatty acids in hepatocytes, alternative pathways of fatty acid oxidation are activated, such as ß-oxidation in peroxisomes. The peroxisomal acylCoA oxidases ACOX is one of the first and rate-limiting enzymes of ß-oxidation pathways in peroxisomes (Reddy and Hashimoto, 2001). Expression of ACOX increase in MASLD patients indicating that peroxisomal ß-oxidation is compensatively enhanced in MASLD (Motoyuki et al., 2007). In our study we observed an increase in ACOX gene expression in PA treated cells, whereas in LS treated cells a significant decrease in its expression was shown. LS reduced the ß-oxidation in peroxisomes indicative of its role in lowering the excessive ROS production which occurs as a result of excessive oxidation of fatty acids. The antioxidant capacity of LS was confirmed through DPPH and ROS assays, where it demonstrated strong radical scavenging activity and suppressed PA-induced oxidative stress. This antioxidant activity likely contributes to its hepatoprotective and anti-inflammatory effects, as evidenced by the downregulation of pro-inflammatory markers TNF-α and ATF6.

Consistent with the in vitro findings, in vivo studies in high-fat diet (HFD)-induced MASLD rats validated the hepatoprotective effects of LS. Treatment with LS significantly prevented body weight gain, reduced liver weight, improved liver histology, and decreased NAFLD Activity Score (NAS). These changes were accompanied by normalization of serum lipid profiles, including reductions in total cholesterol, triglycerides, and VLDL levels, further confirming the lipid-lowering effect of LS.

Beyond the liver, LS demonstrated robust anti-adipogenic activity in 3T3-L1 adipocytes by inhibiting adipocyte differentiation and downregulating the expression of *Pparγ, Srebp-1c, Cebpα* and *Fabp4*. This suggests that LS suppresses lipid accumulation not only in hepatocytes but also in adipose tissue. These anti-adipogenic effects were recapitulated *in vivo*, where LS treatment led to reduced adipose tissue mass and preserved adipocyte morphology, thereby preventing adipose tissue hypertrophy and inflammation. Moreover, LS inhibited pancreatic lipase activity in a dose-dependent manner, suggesting its potential to limit dietary fat absorption, a key factor in obesity and MASLD progression.

Importantly, LS exhibited marked benefits on glucose metabolism. Oral Glucose Tolerance Tests (OGTT) revealed enhanced glucose clearance and reduced blood glucose levels in LS-treated rats. Histological analyses showed restoration of pancreatic islet architecture and β-cell mass, particularly in the high-dose LS group (LSHD). This was paralleled by a significant increase in insulin and GLP-1 secretion, indicating improved pancreatic function and incretin signalling.

## Conclusion

Collectively, our findings establish *Lodhrasavam* as a promising multitargeted therapeutic for MASLD. Its ability to modulate key components of the liver–adipose–pancreas axis—by reducing hepatic steatosis, inhibiting adipogenesis, and enhancing pancreatic endocrine function, underscores its potential to holistically address the metabolic and histological derangements associated with MASLD. These results position LS as a compelling polyherbal candidate for the prevention and management of MASLD and associated metabolic syndromes.

## Acknowledgements

The authors gratefully acknowledge the Indian Council of Medical Research (ICMR) for awarding the ICMR-SRF fellowship to Sania Kouser [File No. 3/1/2(13)/OBS/2022-NCD-II]. The authors acknowledge JM Financial for funding the preclinical studies. The authors also thank the University of Trans-Disciplinary health sciences and technology (TDU), Bangalore, for providing the necessary infrastructure and facilities to carry out the experiments and the Rural India Support Trust (RIST) for financial support. The authors extend their appreciation to Mr. Pranav Banvi and Ms. Soumya Garawadmath for their assistance as interns in the laboratory. The authors also acknowledge Dr. Manjunatha P. Mudagal and Dr. Suresh Janadri for their guidance and support during the preclinical studies.

